# HiCORE: Hi-C analysis for identification of core chromatin loops with higher resolution and reliability

**DOI:** 10.1101/2020.10.01.322024

**Authors:** Hongwoo Lee, Pil Joon Seo

## Abstract

Genome-wide chromosome conformation capture (3C)-based high-throughput sequencing (Hi-C) has enabled identification of genome-wide chromatin loops. Because the Hi-C map with restriction fragment resolution is intrinsically associated with sparsity and stochastic noise, Hi-C data are usually binned at particular intervals; however, the binning method has limited reliability, especially at high resolution. Here, we describe a new method called HiCORE, which provides simple pipelines and algorithms to overcome the limitations of single-layered binning and predict core chromatin regions with 3D physical interactions. In this approach, multiple layers of binning with slightly shifted genome coverage are generated, and interacting bins at each layer are integrated to infer narrower regions of chromatin interactions. HiCORE predicts chromatin looping regions with higher resolution and contributes to the identification of the precise positions of potential genomic elements.

**Author Summary:** The Hi-C analysis has enabled to obtain information on 3D interaction of genomes. While various approaches have been developed for the identification of reliable chromatin loops, binning methods have been limitedly improved. We here developed HiCORE algorithm that generates multiple layers of bin-array and specifies core chromatin regions with 3D interactions. We validated our algorithm and provided advantages over conventional binning method. Overall, HiCORE facilitates to predict chromatin loops with higher resolution and reliability, which is particularly relevant in analysis of small genomes.

## Introduction

Hi-C analysis maps all possible genomic interactions in a manner dependent on three-dimensional (3D) distance. In this method, cross-linked chromatin is digested into fragments by a restriction enzyme [1]. Then, restriction fragments are proximity-ligated to generate a library of chimeric circular DNA. Paired-end sequencing and mapping of reads to a reference genome identifies interacting restriction fragments and measures their interaction frequencies [2]. Unfortunately, despite the high sequencing depth of typical Hi-C experiments, single fragment–resolution contact matrices are too sparse to analyze. Hence, to ensure efficient and reliable data processing, Hi-C data are usually binned with fixed intervals at low resolution (>5 kb) [3-5].

Because the resolution of the Hi-C map is a critical issue for 3D genome conformation studies, multiple studies have focused on improving the resolution of raw Hi-C matrices [6-10]. Although many cutting edge technologies have been implemented, including machine learning-based computational methods, the binning strategy is still an important determinant of Hi-C resolution. Two major types of binning methods have been used in practice: fixed interval binning and fragment unit binning. Fixed interval binning allows intervals to have a regular size (e.g., 5 kb) without considering restriction fragment size, whereas fragment unit binning has an interval defined by the size of restriction fragment(s) [5]. Although fixed interval binning is a practical approach that facilitates efficient analysis of Hi-C data, fixed intervals mismatch restriction fragments of variable sizes, especially at a high resolution, creating bias in chromatin loop detection [5]. Thus, fragment unit binning methods have also been improved in parallel for high-resolution Hi-C analysis. Because single fragment resolution analysis is limited due to contact matrix sparsity, as an alternative method, several restriction fragments are assembled to generate a single bin (here called multi-fragment binning) [6, 11].

Following the binning step, genomic looping regions are identified by various methods, which can be roughly divided into two categories depending on the methodology used for chromatin loop detection. The first category, including HiCCUPS [12] and cLoop [13], primarily identifies chromatin loops located at topologically associating domain (TAD) border regions that are associated with enhancer–promoter interactions [14, 15]. The other category, including Fit-HiC2 [16], HOMER (http://homer.ucsd.edu/homer/), and GOTHiC [17], is well suited for the identification of smaller, local chromatin loops independently of TAD structures [16, 18]. Because the methods of the latter category do not account for a high-order 3D folding structure, they require a more robust post-analysis to determine the reliability of chromatin loop prediction.

In this study, we developed the HiCORE algorithm, which includes an advanced binning method and a reliable loop prediction strategy. HiCORE predicts fine-scale interactions between genomic elements. Independently of the binning method, single-layered binning (conventional method) is insufficient for defining chromatin looping regions, especially at high resolution. To further improve resolution and reliability of detecting 3D chromatin interactions, HiCORE generates multiple layers of bin arrays with different genome coverages. Significantly interacting chromatin regions at each layer are identified [16], and then this information from multiple layers is integrated to draw partially overlapping and narrower chromatin regions with 3D interactions, which represent chromatin loops with higher resolution. Prediction of high-resolution chromatin loops allows the estimation of precise positions of genomic elements. We also validated the reliability of the specified chromatin-interacting regions based on their relative statistical significances.

## Results and Discussion

### Framework of HiCORE

HiCORE generated multiple layers of bin arrays, in which a single bin was defined through serial assembly of fragments (multi-fragment binning) to have a size above a certain threshold. The first layer was generated by forward multi-fragment binning initiated from the starting point (5’-end) of the genome (forward binning), and the second layer was created by reverse multi-fragment binning initiated from the end point (3’-end) of the genome (reverse binning). Additional layers were generated by bidirectional binning from randomly selected positions of the genome (random binning). Bin arrays at each layer could be shifted a bit relative to one another, and thus have different genome coverages (Fig 1). An interaction-frequency matrix of fragment–resolution Hi-C data was assigned to bin arrays at each layer, and bin pairs with statistically significant interactions were identified using the Fit-HiC2 package [16]. Consequently, overlapping regions of interacting bins from multiple layers could be drawn, enabling us to specify reliably interacting chromatin regions at higher resolution (Fig 1).

**Fig 1.**
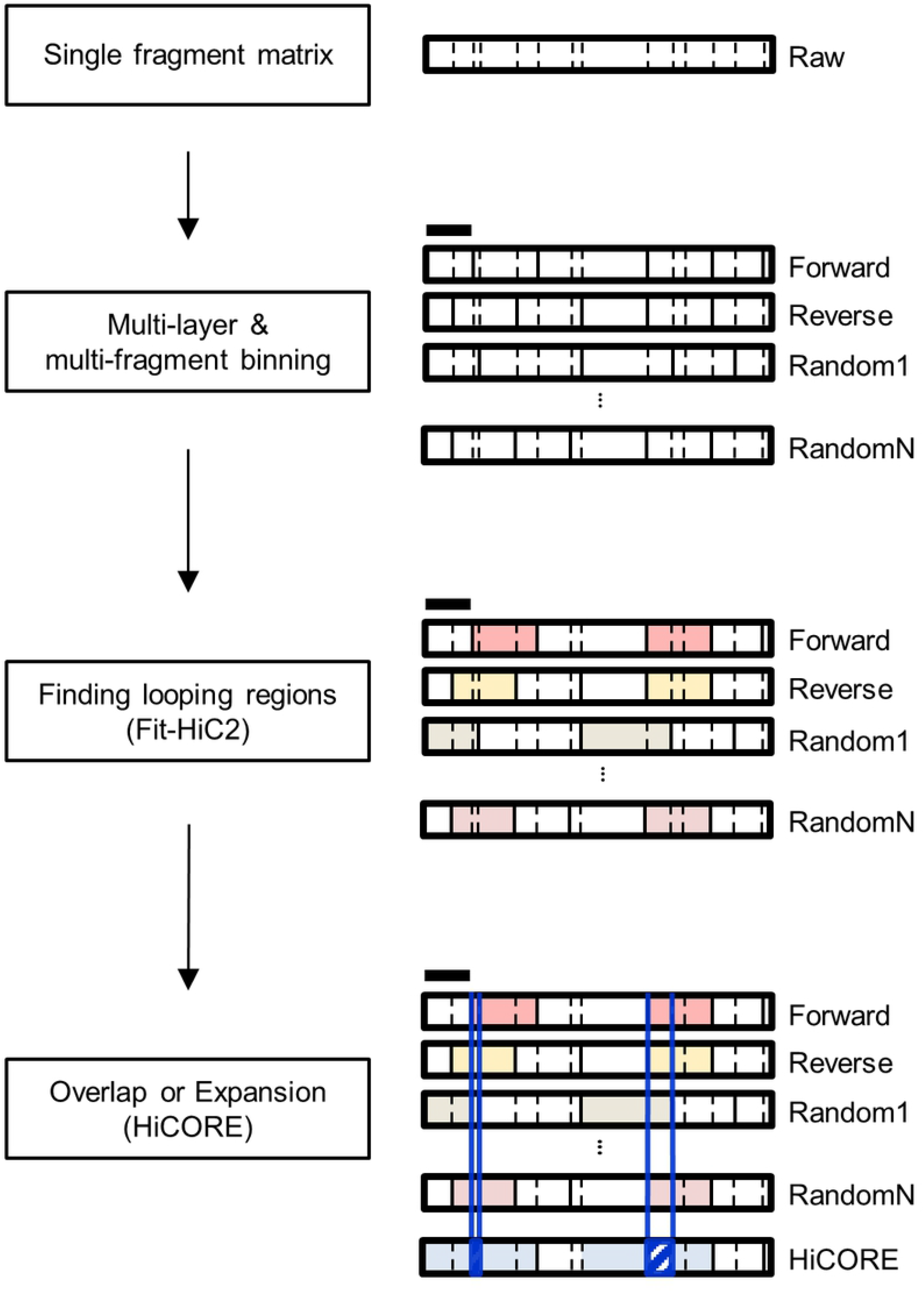
HiCORE features. Schematic diagram showing the working process of HiCORE. Multiple layers of multi-fragment binning are produced, and chromatin looping regions at each layer are predicted based on raw 3D contact matrices. HiCORE identifies overlapped regions of interacting bins from multiple layers, specifying chromatin regions with 3D interaction at higher resolution. Dashed lines indicate restriction sites. Solid lines indicate multi-fragment bin units larger than a given cutoff bin size. Colored regions in each layer indicate chromatin looping regions. HiCORE suggests not only overlapped regions (hatched regions), but also expanded regions (blue-shaded regions).

### Dataset description

As a proof of concept, we used Hi-C data from the model plant *Arabidopsis*, which has a relatively small genome [11]. *Arabidopsis* Hi-C analysis suggested that no clear 3D chromatin conformation exists in the absence of TAD structures [19, 20]. Moreover, genes encoding CCCTC-BINDING FACTOR (CTCF) insulator proteins have not been identified in the *Arabidopsis* genome [21]. However, several studies raised the possibility that chromatin looping is crucial for gene regulation in *Arabidopsis* [11, 22, 23]. Hence, we wanted to implement a stringent method for 3D genome analysis that identifies reliable chromatin looping regions in the model plant genome. We developed a pipeline consisting of HiCORE (for binning) followed by the Fit-HiC2 packages, which can identify the chromatin loops independently of TAD structures [16]. The output of Fit-HiC2 was further post-processed by HiCORE (for loop detection with higher resolution and reliability).

### Increased resolution and reliability in prediction of chromatin loops by HiCORE

To validate the relevance of HiCORE in specifying chromatin looping regions, we generated multiple layers of bin arrays, in which a single bin was defined through multi-fragment binning to have a size above 700bp. The threshold bin size was determined by the raw matrix resolution of the original Hi-C data [14]. We integrated information about 3D chromatin interactions from multiple layers and analyzed the average sizes of specified chromatin regions with 3D contacts in units of fragments. Analysis with a single layer of binning (conventional method) revealed that the average size of all specified chromatin regions with 3D contacts was 4.6 fragments (Fig 2A). However, the HiCORE-generated multiple layers enabled further specification of chromatin looping regions. Compared with single-layered binning, the average size of all specified chromatin regions with 3D contacts was reduced by ∼50% (to 2.2 fragments) after integration of 20 layers (Fig 2A), demonstrating that chromatin looping regions are better specified after HiCORE analysis. Surprisingly, when 20 layers of bin arrays were overlapped, 44% of all predicted chromatin regions with 3D interactions were specified at single-fragment resolution (S1 and S2 Fig). We further tested our HiCORE algorithm with various cutoff values in the threshold bin size from 400 to 2000 bp, and confirmed that chromatin looping regions could be further specified by HiCORE in all threshold size examined (S3 Fig). In addition, the HiCORE-generated loops had higher reliability, which was supported by lower *q*-values in Fit-HiC2 analysis (Fig 2B).

**Fig 2.**
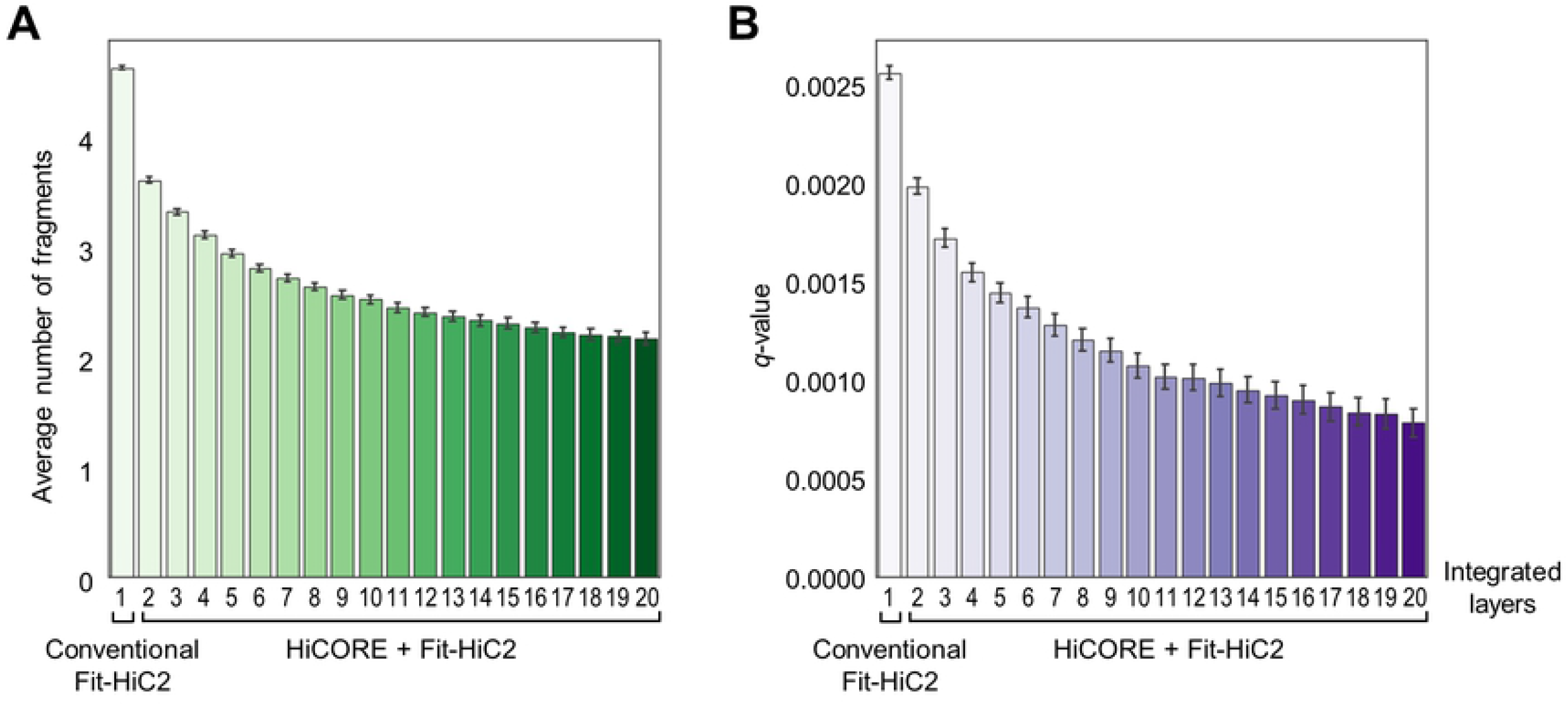
HiCORE test results. (**A**) Average number of fragments in each specified chromatin-interacting region. (**B**) Average *q*-values (minimum false discovery rate, FDR) of specified interacting chromatin regions. The *q*-values were estimated by Fit-HiC2 packages [16]. In (**A**) and (**B**), for binning at each layer, restriction fragments smaller than 700 bp are merged with neighboring fragments until the bin is larger than 700 bp. One bin layer was added at a time, repeatedly, and chromatin looping regions were specified. The x-axis indicates the number of layers integrated for HiCORE analysis.

In support, chromatin looping regions were better specified throughout the genome, compared with conventional single-layered Fit-HiC2 analysis. For example, while three chromatin loops were predicted by the conventional Fit-HiC2 analysis within Chr1:590 kb – 599 kb regions (Fig 3A), HiCORE selected a reliable chromatin loop (shown in blue) and defined the chromatin looping region with higher resolution (Fig 3B): one side specified at a single-fragment unit and the other at a 2-fragment unit.

**Fig 3.**
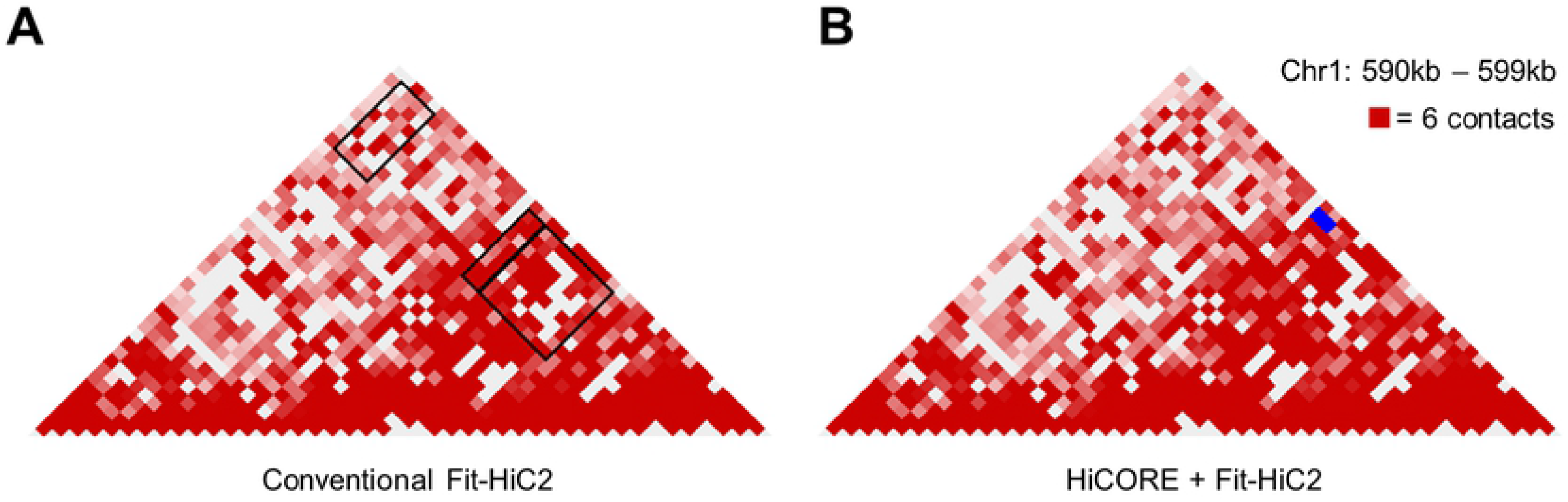
Comparison between conventional and HiCORE analysis in specification of chromatin looping regions. (**A**) Chromatin loops predicted by conventional binning method. Chromatin regions with 3D interactions are indicated by black boxes. (**B**) Chromatin loops predicted by HiCORE analysis. Chromatin regions with 3D interactions are indicated by blue boxes. The interaction frequency was shown at single fragment scale.

Notably, HiCORE results could not be obtained by single-layered single fragment binning. HiCORE analysis predicted a total of 1987 chromatin loops, 27% of which (541 chromatin loops) were specified at single fragment resolution (S4 Fig). However, the single-layered Fit-HiC2 method identified 185 chromatin loops at single fragment resolution (Fig 4A). Furthermore, the average size of HiCORE-specified chromatin regions with 3D contacts (for 1987 chromatin loops) was much smaller than that by the conventional single-layered Fit-HiC2 analysis (for 185 chromatin loops) (Fig 4B). The chromatin loops predicted by HiCORE had also lower *q*-value (Fig 4A), indicating that HiCORE enables the successful identification of chromatin looping regions at higher resolution.

**Fig 4.**
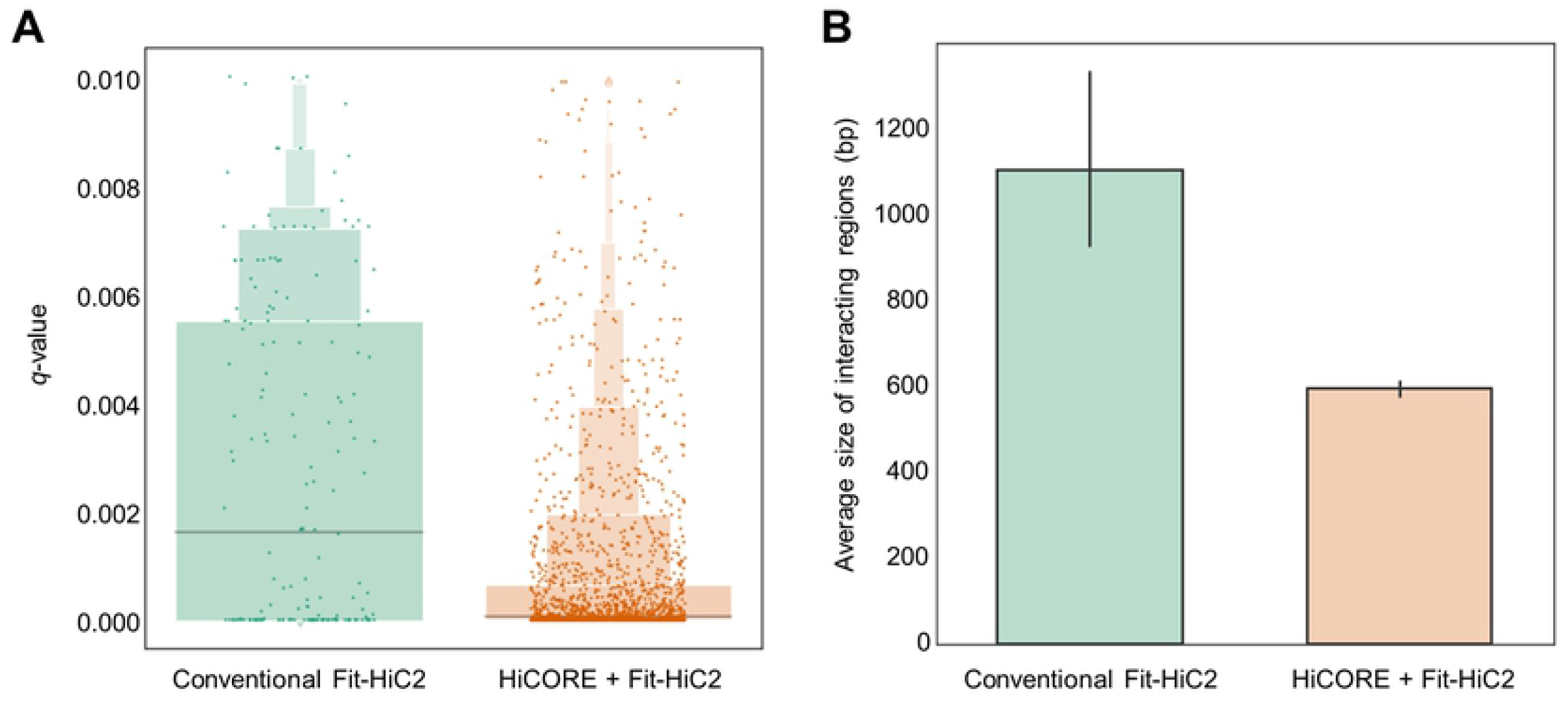
Comparison between conventional single-fragment binning method and HiCORE analysis. (**A**) The box plot of *q*-value distribution. (**B**) Average size (bp) of specified interacting chromatin region. For HiCORE analysis, restriction fragments smaller than 700bp are merged with neighboring fragments until the bin is larger than 700bp. Twenty bin layers were integrated, and chromatin looping regions were specified.

We also showed the results in units of base-pair length (S5 Fig). However, at extremely high resolution (cutoff value < matrix resolution), the average size of all specified chromatin regions with 3D contacts in units of base-pair length did not gradually decrease as binning layers were integrated (S5 Fig), although the average fragment number was reduced (Fig 2A). The inconsistency was possibly due to the sparsity of 3D contact matrices in smaller bins, which caused low reproducibility at each layer and thereby excluded during HiCORE analysis. Thus, HiCORE can specify the core chromatin looping regions, but optimal analysis can usually be performed above the matrix resolution.

### Additional applications of HiCORE

The HiCORE method can also be used to profile all possible chromatin loop information predicted by multiple binning layers. The single-layered binning can introduce significant bias into the identification of chromatin loops. For instance, a chromatin loop in the *Arabidopsis FLC* locus was empirically validated [11, 23], but this loop was predicted by only a few binning layers (Fig 5). This observation indicates that there is no best single binning layer that sufficiently covers genomic interactions, and that comprehensive integration of chromatin looping information predicted by multi-layered binning with different genome coverage is required. HiCORE can provide a platform for data integration and optionally suggest a wide range for possible chromatin looping regions (Fig 1). Furthermore, using HiCORE, chromatin looping regions predicted by only a few layers could be selectively integrated to specify narrower overlapping chromatin regions. Indeed, the empirically-validated *FLC* loop was precisely specified by HiCORE analysis with integration of selected binning layers that had statistically significant 3D-interaction values within the *FLC* locus (Fig 5).

**Fig 5.**
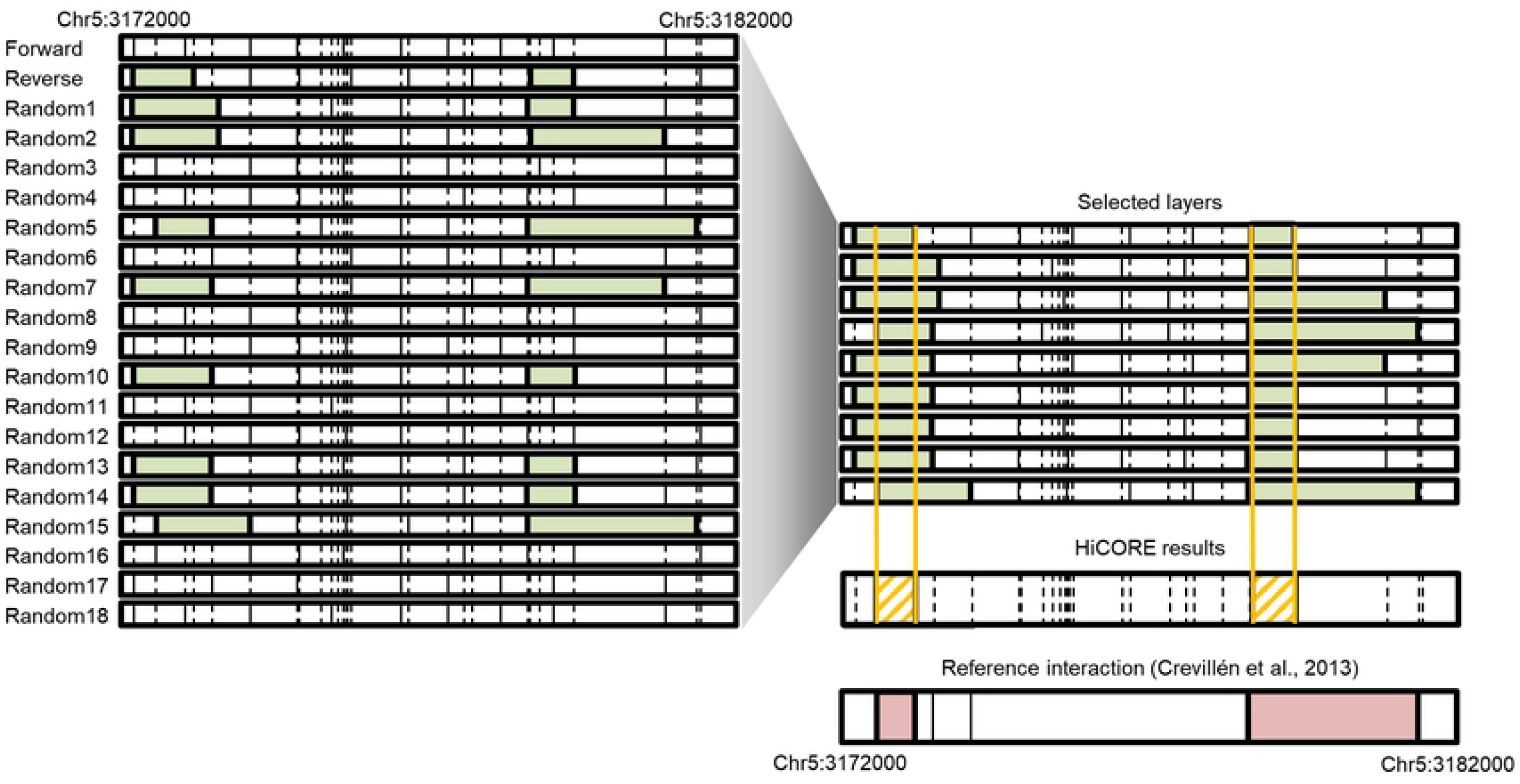
Chromatin loops within the *FLC* locus. For binning at each layer, restriction fragments smaller than 700bp were merged with neighboring fragments until the bin is larger than 700bp. Twenty layers of binning were produced. For each binning layer, chromatin loops in the genomic coordinates between (<Chr5:3179000-3181500> : <Chr5:3172000-3174000>) are predicted (Green). Then, bin layers with the *FLC* loop prediction were selected, and the overlapped regions were specified by HiCORE (Yellow). The empirically-proven interaction around the *FLC* locus (<Chr5:3172571-3173171> : <Chr5:3178619-3181368> is shown in red.

Hi-C is a commonly used technique for mapping 3D chromatin organization. Despite significant advances in the accuracy and resolution of genome contacts, identification of fine-scale chromatin loops remains challenging, mostly because of low sequencing depth and single-layered binning. The HiCORE algorithm takes advantage of an advanced binning method, and also predicts higher-resolution chromatin looping by generating multiple layers of bin arrays, which are slightly shifted relative to each other. Extraction of partially overlapping, more narrowly interacting regions allows higher-resolution analysis of chromatin looping from Hi-C data. HiCORE can be incorporated into current Hi-C data analysis pipelines, improving the resolution in predicting chromatin looping regions. HiCORE analysis will facilitate better separation of genomic contacts and the identification of potential genomic elements.

## Method

### Hi-C data and their pre-processing

The high resolution Hi-C data of *Arabidopsis* were obtained from NCBI Sequence Read Archive (SRA; http://www.ncbi.nlm.gov/sra/) under accession number SRP064711. The 4 replicates were mapped to *Arabidopsis* genome (TAIR10) and further processed by Juicer packages [12]. We extracted the raw interaction matrix from the ‘.hic’ file that was derived from reads with mapping quality > 30.

### Binning strategy

Fragments with a size below a given cutoff length are merged with neighboring fragments. The merged fragments are assembled into a single bin (multi-fragment bin). This binning strategy is applied to all fragments except for the end fragment. For ‘forward binning’, from the 5’ end of the chromosome, a restriction fragment shorter than the cutoff length is merged with the following fragment (in the 3’-direction) until the merged fragment is longer than cutoff length. For ‘reverse binning’, from the 3’ end of the chromosome, a restriction fragment shorter than the cutoff length is merged with the preceding fragment (in the 5’-direction) until the merged fragment is longer than the cutoff length. For ‘random binning’, HiCORE randomly selects restriction fragments for construction of a binning layer. Randomly selected fragments shorter than the cutoff length are first merged with the following fragment in the 3’-direction. In the middle of fragment assembly, if the next fragment in the 3’-position was already merged with other fragments, the bin being generated is merged with the fragment in the 5’-direction until the bin being generated is longer than the cutoff length. If the neighboring fragments in both directions are already taken by other completed bins, the bin being generated is divided into two units. The first unit (5’-neighboring fragments of the random fragment) and the second unit (the random fragment plus its 3’-neighboring fragments) of the bin being generated are separately merged to already-completed neighboring bins in the 5’ and 3’ positions, respectively.

### Single-layered Fit-HiC2 analysis

Using the extracted raw fragment matrix file, we generated ‘.interaction.gz’ and ‘fragments.gz’ files that are necessary for subsequent Fit-HiC2 analysis. We normalized the raw matrix using ‘HiCKRy.py’ in Fit-HiC2 packages [16]. Fit-HiC2 analysis was performed with the following parameters : -r 0 -L 2000 -U 25000 -x intraOnly. The output loops were filtered by a statistical cutoff value (*q* < 0.01)

### Profiling of core chromatin looping regions using HiCORE

The raw fragment matrix file was assigned to each bin layer, and then single-layered Fit-HiC2 analysis was performed in each bin layer. After defining the interacting chromatin regions using Fit-HiC2 packages, a statistical cutoff value (*q* < 0.01) is used to obtain statistically significant chromatin regions with 3D interactions. Fit-HiC2 output files, which contain 3D interaction data in units of multi-fragment bins, are converted to data in units of single-fragment bins for each binning layer. Then, the output files are further processed by HiCORE to provide either overlapped core interacting chromatin regions or expanded interacting chromatin regions using single-fragment data from multiple binning layers. The HiCORE-processed data of single-fragment interactions are shown in units of the original multi-fragment bins for comparison of specification of chromatin looping regions.

### Software availability

HiCORE is implemented as a Python3 package. The most current stable version is available at: https://github.com/CDL-HongwooLee/HiCORE

## Acknowledgements

We thank Dr. Sangrea Shim (Seoul National University, Korea) for fruitful discussion.

## Funding

This work was supported by the Basic Science Research (NRF-2019R1A2C2006915) and Basic Research Laboratory (2020R1A4A2002901) programs provided by the National Research Foundation of Korea and by the Next-Generation BioGreen 21 Program (PJ01314501) provided by the Rural Development Administration.

## Conflict of Interest

none declared.

## Supporting information

**S1 Text**. The information for HiCORE implementation.

**Supplemental Figure 1. Box plot for average number of fragments in each specified chromatin-interacting region**.

Restriction fragments smaller than the cutoff values (700bp) were merged with neighboring fragments until the bin is larger than the cutoff size. One bin layer was added at a time, repeatedly, and chromatin looping regions were specified. The x-axis indicates the number of layers integrated for HiCORE analysis. See also Figure 2A.

**Supplemental Figure 2. Size distribution of HiCORE-specified looping regions**.

Restriction fragments smaller than the cutoff values (700bp) were merged with neighboring fragments until the bin is larger than the cutoff size. Twenty bin layers were integrated by HiCORE and the overlapped looping regions were identified. Specified regions in each side of a chromatin loop was counted independently. The number of interacting regions was shown based on the number of consisting fragment(s) in a HiCORE-specified region.

**Supplemental Figure 3. The average number of fragments in each specified interacting chromatin region, with various cutoff values**.

Restriction fragments smaller than the cutoff values were merged with neighboring fragments until the bin is larger than the cutoff size. One bin layer was added at a time, repeatedly, and chromatin looping regions were specified. The x-axis indicates the number of layers integrated for HiCORE analysis.

**Supplemental Figure 4. The fragment number distribution of HiCORE-specified chromatin loops**.

Restriction fragments smaller than the cutoff values (700bp) were merged with neighboring fragments until the bin is larger than the cutoff size. Twenty bin layers were integrated by HiCORE, and the overlapped looping regions were identified. Reliable chromatin loops were collected, and they were categorized based on the number of consisting fragment(s) in a pair of HiCORE-specified chromatin looping regions. 1F, 1-fragment unit; 2F, 1∼2-fragment unit; 3F, 1∼3-fragment unit; 4F, 1∼4-fragment unit; >4F, >4-fragment unit.

**Supplemental Figure 5. The average size of each specified interacting chromatin region**.

Restriction fragments smaller than the cutoff values were merged with neighboring fragments until the bin is larger than the cutoff size. One bin layer was added at a time, repeatedly, and chromatin looping regions were specified. The x-axis indicates the number of layers integrated for HiCORE analysis.

